# Isolation and identification of enteroviruses from sewage and sewage-contaminated water samples from Ibadan, Nigeria, 2012-2013

**DOI:** 10.1101/189902

**Authors:** Johnson Adekunle Adeniji, Adetunji Oladapo Adewale, Temitope Oluwasegun Cephas Faleye, Moses Olubusuyi Adewumi

## Abstract

In 2010, we described sewage contaminated water (SCW) bodies that consistently yielded enteroviruses (EVs) in enterovirus surveillance (ES) sites in Lagos, Nigeria. By 2012, we demonstrated the presence and circulation of Wild Poliovirus 3 (WPV3) in these ES sites. Here we describe ES sites that consistently yield EVs in Ibadan metropolis southwest Nigeria.

Twenty-five ES samples were collected by grab method from nine sites between October, 2012 and March, 2013. Samples were concentrated and four (RD, HEp2C, MCF-7 and L20B) different cell lines used for virus isolation from the concentrates. Isolates were subjected to RNA extraction, cDNA synthesis, PanEnterovirus 5^*l*^-UTR and VP1 assays. Unidentifiable isolates were further subjected to species-specific RT-PCR assays. Amplicons were sequenced, isolates identified and subjected to phylogenetic analysis.

Twenty-five isolates were recovered from 8 (32%) of the 25 ES samples collected. Twenty-three of the isolates were identified as EVs by the PanEntero5^*l*^-UTR assay. Thirteen (57%) of the 23 EVs were positive for the VP1 assay, and identified as Coxsackievirus B3 (CVB3) (1 isolate), CVB6 (1 isolate), E6 (2 isolates), E7 (5 isolates), E11 (1 isolate), E12 (1 isolate) and E13 (2 isolates). None and 2 (25%) of the remaining isolates were positive for the EV-B and EV-C assays, respectively. The 2 EV-C positive enteroviruses were isolated on MCF-7.

This study describes three very productive ES sites, and documents the presence of CVB3, CVB6, E6, E7, E11, E12 and, E13 in Ibadan, Nigeria. It shows that including other cell lines in EV isolation protocols can broaden the diversity of EV types recoverable.

## INTRODUCTION

Enteroviruses (EVs) are members of the family *Picornaviridae*, order *Picornavirales*. Within the genus are 13 species. The best studied member is poliovirus (PV) and is a member of EV Species C (EV-C), while EV-B has the most members (over 60) (www.picornaviridae.com). An EV virion is an icosahedron with a diameter of 27 – 30nm. The naked capsid encloses a 5^*l*^ protein-linked, single-stranded, positive-sense, ~7.5kb, RNA genome. The single open reading frame (ORF), encoded by the genome, is flanked by 5^*l*^ and 3^*l*^ untranslated regions (UTRs). The polyprotein product of the ORF is initially auto-catalytically cleaved into three smaller proteins (P1, P2 and P3) and ultimately into 11 proteins (VP1-VP4 from P1, 2A-2C from P2 & 3A-3D from P3). An association was demonstrated between the nucleotide and amino acid sequence of VP1 and EV types (Oberste*et al*., 1999, 2000). Consequently, VP1 sequence data is now used for EV identification (Oberste*et al*., 2000,2003; Casas *et al*., 2001; Norderet*al*., 2001; Caro *et al*., 2001; Blomqvist*et al*., 2008, Adeniji and Faleye 2014a).

The Global Polio Eradication Initiative (GPEI) was established with the goal of eradicating poliovirus (WHO, 1988). It uses vaccination (Oral Polio Vaccine [OPV], Inactivated Polio Vaccine [IPV]) and surveillance as the major tools to achieve its purpose. Surveillance is done using case based (CBS) and environmental (ES) surveillance. While CBS (as implemented in acute flaccid paralysis [AFP] surveillance) uses clinical manifestations as the sign post for finding the virus (WHO, 2004), ES looks for the virus in sewage and/or sewage contaminated water (SCW) (WHO, 2003). The ES strategy is predicated on the finding that, though not all individuals infected with EVs show clinical manifestations, all shed the virion in stool in large amounts for weeks (Ranta*et al*., 2001), and consequently into sewage and SCW (Ranta*et al*.,2001). Hence, sewage and SCW can serve as a good source for finding virus circulating in a population without or before clinical manifestations show up (WHO, 2003).

In 2010, we (Adeniji and Faleye, 2014a) described ES sites in Lagos, Nigeria that consistently yielded enteroviruses (EVs). By 2012, we (Faleye and Adeniji, 2015a) demonstrated the presence and circulation of Wild Poliovirus 3 (WPV3) in these ES sites. These discoveries were followed by immunization campaigns and till date those findings mark the last sightings of WPV3 globally (Asghar et al., 2014). We subsequently handed over these sampling sites to the WHO poliovirus ES program in Nigeria. Against this backdrop, we extended surveillance to Ibadan metropolis (the largest city [by land mass] in West-Africa) in Southwest Nigeria, a place that was not and is still currently not being sampled by the WHO poliovirus Environmental Surveillance program. As we did in Lagos (Adeniji and Faleye, 2014a, Faleye and Adeniji, 2015a & b), here we describe ES sites that consistently yielded enteroviruses in Ibadan, Southwest, Nigeria.

## MATERIALS AND METHOD

### Sample Collection and Concentration

Sewage and sewage-contaminated water (SCW) samples were collected from nine (9) selected sites in Ibadan metropolis, South-western Nigeria. Sampling was of convenience and most sites were sampled repeatedly (at least once every two months) between October 2012 and March 2013. At every sampling, one liter of sewage or SCW was collected into clean plastic bottles before 9:00 am using the grab method. The samples were transported in an ice chest to the laboratory in the Department of Virology, College of Medicine, University of Ibadan, Nigeria. In all, 25 samples were collected during this study.

To concentrate the virus particles in the sample, 480mL of each sample was aliquoted in 40 mL volumes into twelve 50 mL centrifuge tubes and first clarified by centrifugation at 3,000 rpm for 1 hour at 4^0^C. The supernatant was pulled into a beaker to which 30g (6%) of polyethylene glycol (PEG) 6000 and 13g (2.6%) of NaCl were added and stirred for 20 minutes (Killington et al., 1996). Subsequently, the mixture was poured into a separation funnel and refrigerated at 4^°^C overnight. The pellets were also pulled and refrigerated overnight.

By the following morning, about 10mL of the bottom layer of the mixture was gently collected into the pooled pellet, and chloroform was added up to 10% of its volume. This was vortexed for 1 minute and subsequently centrifuged at 1,500 rpm for 20 minutes at 4^0^C. Afterwards, the supernatant (subsequently called the concentrate) was gently collected and stored at ‐20^0^C till used for virus isolation. Usually virus isolation commenced the same day.

### Virus Isolation

Four different cell lines were used for virus isolation in this study. These include RD (human rhabdomyosarcoma), Hep-2C (human laryngeal carcinoma), MCF-7 (human mammary gland adenocarcinoma) and L20B (genetically engineered mouse cell line, expressing the human poliovirus receptor CD155) cell lines. All were maintained in medium supplemented with 2% fetal bovine serum (FBS). Cell lines were supplied by the WHO Polio Laboratory, Department of Virology, University of Ibadan, Oyo State, Nigeria.

Each concentrate was inoculated into eight cell culture tubes of each cell line (RD, Hep2C, MCF-7 and L20B) containing a monolayer of cells. Precisely, 200μL of each concentrate was inoculated into each cell culture tube (i.e. 1.6 mL per cell line), and ultimately, 6.4mL of concentrate per sample was analyzed. Tubes were then incubated at 37^°^C and checked for cytopathic effect (CPE; cell rounding, refractiveness and detachment from culture surface) every 24 hours over a period of 7 days. Cells with CPE were suspected to be positive, freeze-thawed thrice and passaged in fresh cultures of the same cell line. Isolates confirmed were kept at ‐20^°^C for further analysis.

Cell culture tubes that developed no CPE by the end of the first 7days of observation, were freeze-thawed three times and passaged in fresh cultures of the same cell line for another 7 days. Hence, negative cultures were examined for a total of at least 14 days before being discarded. Throughout the study period, two tubes of each cell line were labeled as negative control and kept under the same conditions as the experiments. The algorithm followed in this study is depicted in Figure 1.

**Figure 1:**
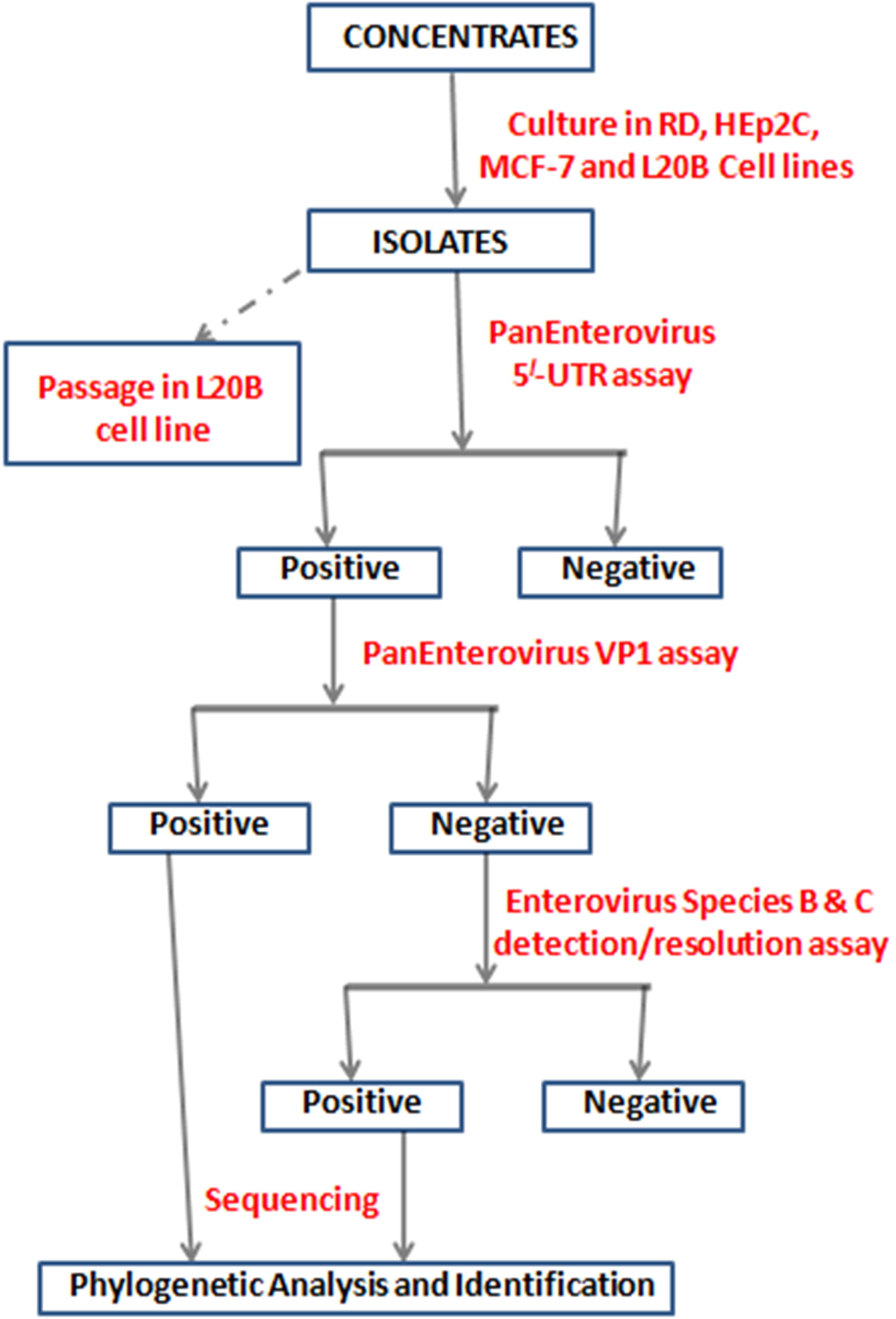
A schematic representation of the algorithm followed in this study.

### Cell culture screening of isolates for Poliovirus

All isolates were passaged in L20B cell line and examined for 5 days to exclude the possibility that recovered isolates were poliovirus. The appearance of CPE in L20B cell line within 24 to 48 hours confirmed the likely presence of poliovirus.

### RNA extraction and cDNA synthesis

All isolates were subjected to RNA extraction using an RNA extraction kit (JenaBioscience, Jena, Germany) in accordance with the manufacturer’s instructions. Similarly, cDNA synthesis was done using Script cDNA Synthesis Kit (Jena Bioscience, Jena, Germany) in accordance with the manufacturer’s instructions. Particularly, cDNA was done using random hexamers as previously discussed (Adeniji and Faleye, 2014b).

### Polymerase Chain Reaction (PCR)

As shown in the algorithm (Figure 1), four PCR assays were run. One each of 5′-UTR (Oberste et al., 2005), VP1 (Oberste et al., 2003), EV-B (Bailly et al., 2011) and EV-C (Adeniji and Faleye, 2014b) PCR assays. The 5′-UTR assay detects all enteroviruses present but cannot be used to determine enterovirus types. The sequence data generated for the VP1 assay on the other hand is sufficient, and used to determine enterovirus types. The EV-B and EV-C PCR assays are species detection/resolution assays targeting the P3 nonstructural region of the genome (Bailly et al., 2011; Adeniji and Faleye, 2014b).

All isolates were subjected to the 5′-UTR assay. Those positive for the 5′-UTR assay were subjected to the VP1 assay. Isolates negative for the VP1 assay were subjected to both the EV-B and EV-C PCR assays. All PCR assays were carried out in 25μL reaction volumes. Primers were made in 100 μM concentrations. Each 25μL reaction contained 5μL of Red Load Taq (Jena Bioscience), 3μL of cDNA, 0.25μL of each primer and 16.5μL of RNase-free water. Thermal cycling was done as follows: 94^0^C for 3 minutes, 45 cycles at 94^0^C for 30 seconds, 42^0^C for 30 seconds and 60^0^C for 30 seconds, with 40% ramp from 42^0^C to 60^0^C which was followed by 72^0^C for 7 minutes and held at 4^0^C till terminated.

It is important to note that the extension time varied based on the length of expected amplicon (Table 1). The reaction products were analysed by electrophoresis in a 2% agarose gel, stained with ethidium bromide.

**Table 1:** The PCR assays used in this study, expected amplicon size and extension time.

| **ASSAY** | **AMPLICON SIZE** | **PCR EXTENSION TIME** |
| --- | --- | --- |
| PanEntero 5'-UTR | 114bp | 30 seconds |
| PanEntero VP1 | 350bp | 30 seconds |
| EV-B Screen | 1,140bp | 90 seconds |
| EV-C Screen | 420bp | 30 seconds |

### Amplicon sequencing and enterovirus typing

Amplicons positive for the VP1 and EV-C PCR assays were shipped to Macrogen Inc, Seoul, South Korea, for purification and sequencing using the respective forward and reverse primers. The Enterovirus Genotyping Tool (EGT) was subsequently used for enterovirus genotyping and species determination (Kroneman et al., 2011).

### Phylogenetic Analysis

The CLUSTAL W program in the MEGA 5 software (Tamura et al., 2011) was used, with default settings, to align sequences of the EV strains recovered in this study with those of reference strains retrieved from GenBank. Using the MEGA 5 software (Tamura et al., 2011), phylogenetic (neighbor-joining) trees were subsequently constructed using the Kimura-2 parameter model (Kimura, 1980) and 1,000 bootstrap replicates.

### Nucleotide sequence accession numbers

All sequencesreported in this study have been deposited in GenBank. Their accession numbers are MF925345-MF925357 and MF957224-MF957225 for the VP1 and 3D sequences, respectively.

## RESULTS

### Sample Collection and Virus Isolation

Twenty-five samples were analyzed in this study. A total of 25 isolates were recovered from the 25 samples analyzed. All the isolates were recovered from eight (32.0%) of the samples collected. All the eight samples were collected from three (33.3%) of the nine sites sampled in this study. The remaining 17 (68.0%) samples did not yield isolates evidenced by the development of CPE (Table 2). With respect to the cell lines used for virus isolation, RD, Hep2C, MCF and L20B yielded 16 (64%), 4 (16%), 5 (20%) and zero (0%) isolates respectively (Table 3). When passaged on L20B cell line, none of the 25 isolates produced CPE within 48 hours, and even up to five days.

**Table 2:**
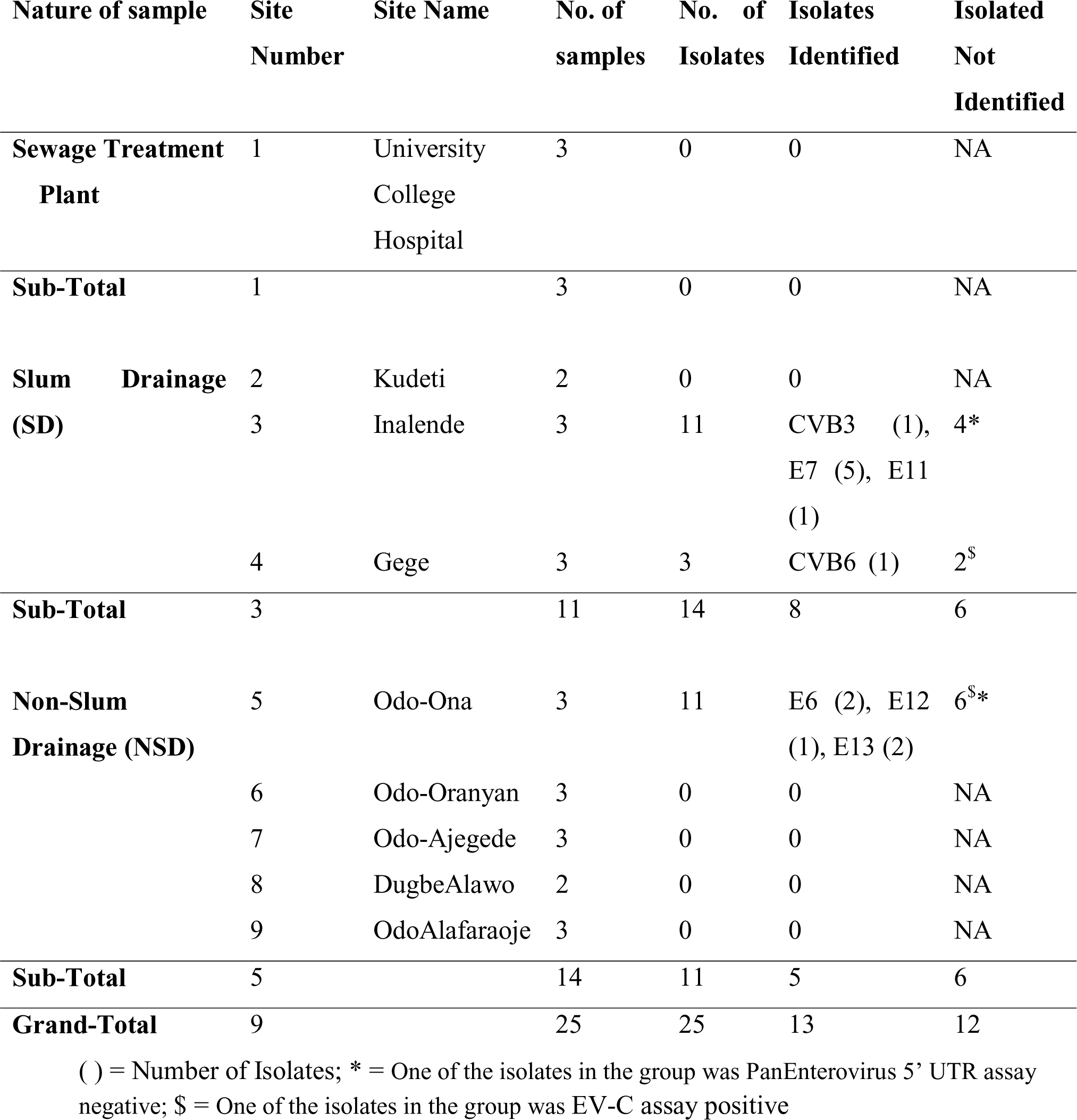
Enterovirus isolation results and types recovered.

**Table 3:**
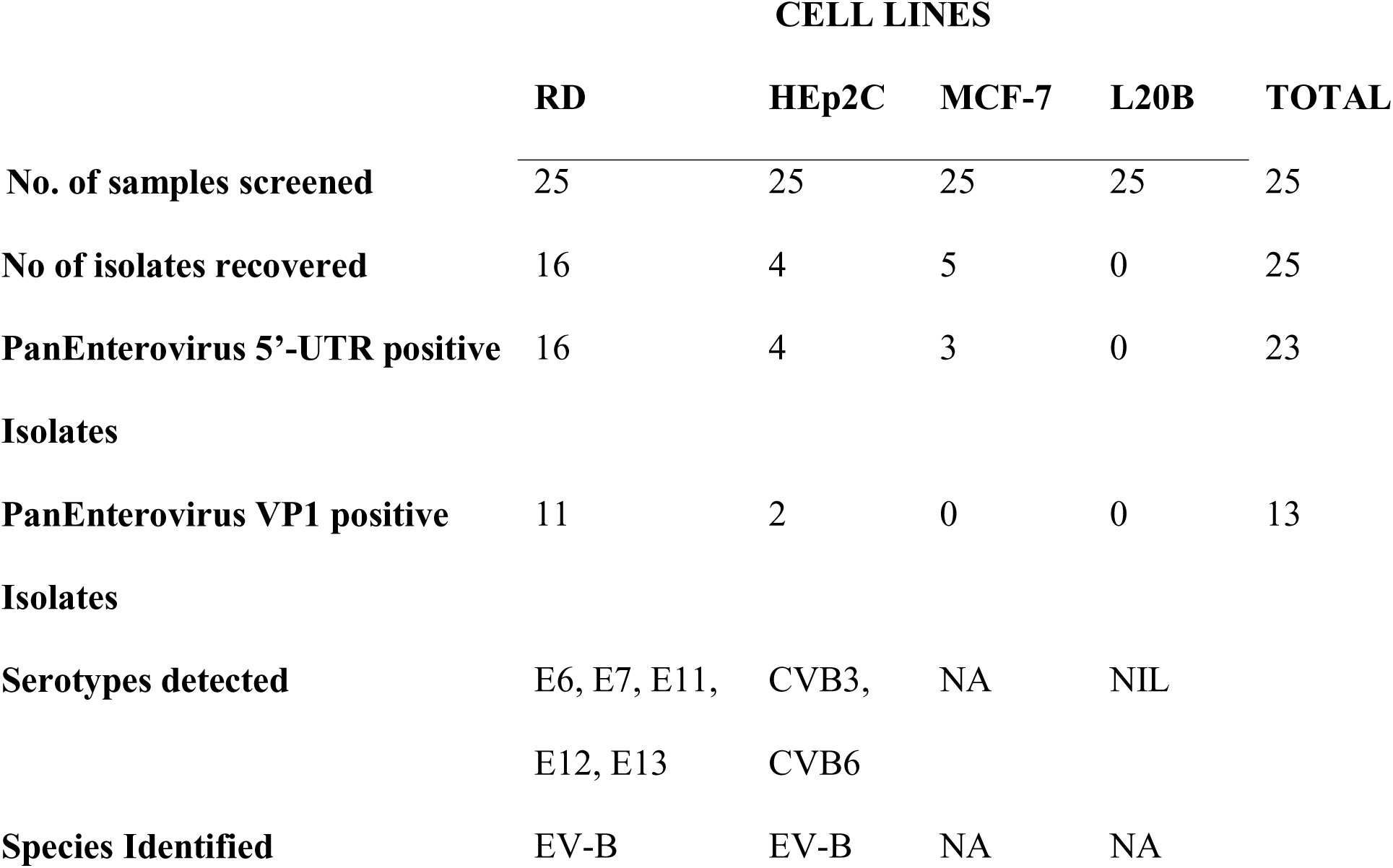
Characterization of Isolates.

### Polymerase Chain Reaction (PCR)

The required 114bp amplicon for the PanEnterovirus 5’-UTR assay was successfully amplified in 23 (92%) of the 25 isolates recovered (Table 3). Hence, the 23 isolates were considered positive for the PanEnterovirus 5’-UTR assay. The remaining two isolates that were negative for this assay were both isolated on MCF-7 cell line (Table 3). The two isolates were recovered from sample sites 3 and 5 (Table 2).

The required ~350bp amplicon for the PanEnterovirus VP1 assay was successfully amplified in 13 (56.5%) of the 23 isolates positive for the PanEnterovirus 5’-UTR assay (Table 3). Hence, 13 and 10 (43.5%) isolates were considered positive and negative, respectively, for the PanEnterovirus VP1 assay. The 10 isolates negative for the PanEnterovirus VP1 assay were isolated on RD (5 isolates), HEp2C (2 isolates) and MCF-7 (3 isolates) cell lines (Table 3).

The 10 enterovirus isolates that were negative for the PanEnterovirus VP1 assay were subjected to both EV-B and EV-C PCR assays. All 10 isolates were negative for the EV-B assay (Table 4). On the other hand, 2 of these isolates were positive for the EV-C assay (Table 4). These two (2) isolates were isolated on MCF-7 (Tables 3 & 4), recovered from sample sites 4 and 5 (Table 2), and did not replicate in L20B when passaged in it (Table 3).

**Table 4:** Results of Enterovirus Species detection/resolution assay.

|  | CELL LINES |  |  | TOTAL |
| --- | --- | --- | --- | --- |
|  | RD | HEp2C | MCF-7 |  |
| **No. of EV Isolates completely identified by previous screens** | 5 | 2 | 3 | 10 |
| **EV-B Species screen positive isolates** | 0 | 0 | 0 | 0 |
| **EV-C Species screen positive isolates** | 0 | 0 | 2 | 2 |
| **Isolates that replicated in L20B** | 0 | 0 | 0 | 0 |
| **Species Identified** | NA | NA | EV-C |  |
| **Unidentified EV Isolates** | 5 | 2 | 1 | 8 |

### Enterovirus typing and Phylogenetic Analysis

Using the sequence data generated from the 13 PanEnterovirus VP1 assay positive isolates, the EGT identified them as Coxsakievirus B3 (CVB3) (1 isolate), CVB6 (1 isolate), Echovirus 6 (E6) (2 isolates), E7 (5 isolates), E11 (1isolate), E12 (1 isolate) and E13 (2 isolates) (Table 2). The identity of the 13 isolates was confirmed as such by phylogenetic analysis (Figure 2).

**Figure 2:**
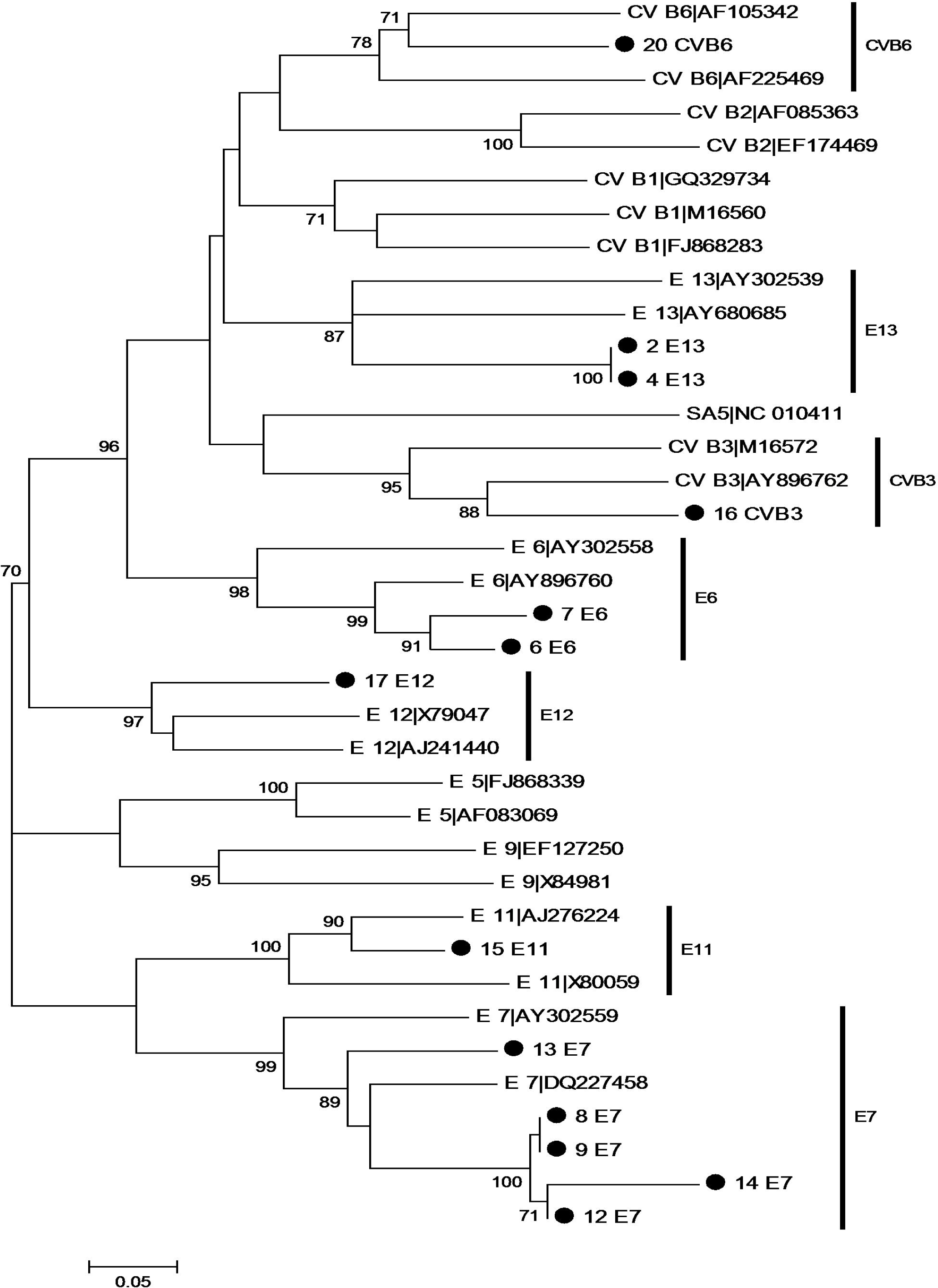
Phylogenetic tree showing all enterovirus isolates recovered from this study alongside reference strains downloaded from GenBank. The tree is based on an alignment of partial VP1 sequences. The newly sequenced strains are highlighted with black circles. The GenBank accession numbers of the reference strains are indicated in the tree. Bootstrap values are indicated if ≥70%.

Considering E7 was the most isolated (5 isolates), it was further subjected to type specific phylogenetic analysis. Two different E7 genotypes were detected in this study; Sub-cluster Nigeria 2 within the sub-Saharan Africa specific cluster and the sub-cluster Nigeria within the Global Cluster (Figure 3). All four isolates in sub-cluster Nigeria 2 (within the sub-Saharan Africa specific cluster) are being described for the first time to the best of the authors’ knowledge. On the other hand, strains closely related to the isolate in sub-cluster Nigeria (within the Global Cluster) were subsequently detected in 2014 in children with AFP in Nigeria (Figure 3).

**Figure 3:**
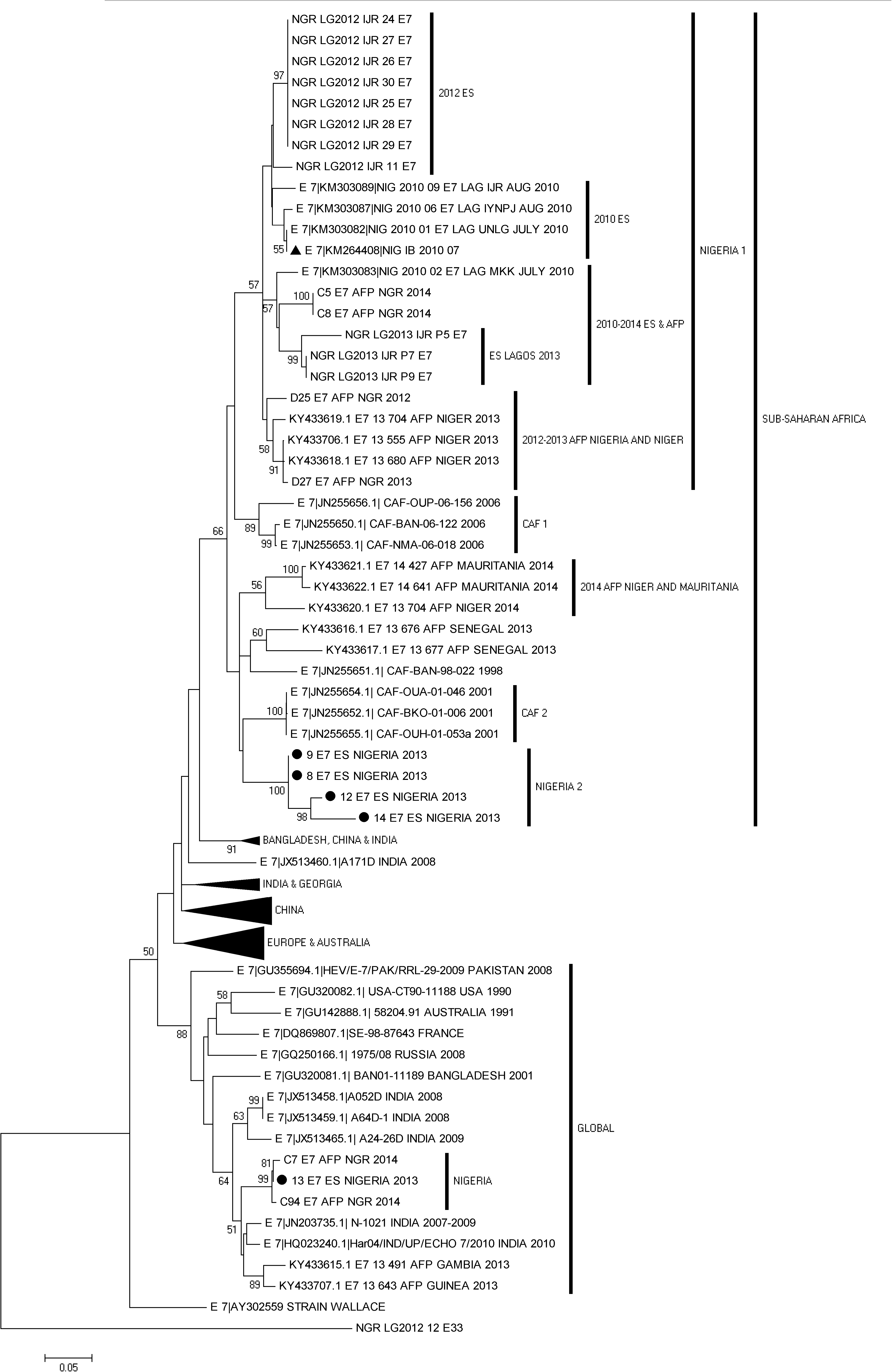
Phylogram of genetic relationship between VP1 nucleotide sequences of E7 isolates. The phylogenetic tree is based on an alignment of the partial VP1 sequences. The newly sequenced strains are indicated with black circles while an E7 strain recovered from the same region in 2010 is indicated with a black triangle. The GenBank accession numbers and strain of the isolates are indicated in the tree if known. Bootstrap values are indicated if >50%.

Using the sequence data generated from the 2 EV-C assay positive isolates, the EGT identified them as truly EV-C. Phylogenetic analysis also confirmed this and further showed that some of the sequences were indigenous to the region. For example, TJ2 shared a common ancestor with the partial 3D region of some recombinant PV2 sequences recovered in Nigeria in 2012. In similar light, TJ1 shared a common ancestor with the partial 3D region of some recombinant PV1 sequences recovered in Cameroon in 2014.

## DISCUSSION

Of the 25 samples screened in this study, EVs were recovered from 32% (8/25). This detection rate is above the 30% threshold stipulated by the WHO (2003). These results, therefore documents the efficiency of the PEG dependent concentration protocol (Killington et al., 1996) used in this study and thus demonstrates its use in enterovirus surveillance. Though, different concentration protocols were used, the detection rate documented here is similar to the 34.6 % we found in Lagos, Nigeria in 2010 when we were scouting for adequate ES sampling sites (Adeniji and Faleye 2014a). By 2012, our EV detection rate in Lagos, Nigeria had risen to 60% (Faleye and Adeniji 2015a) because we had concentrated on sampling sites that repeatedly yielded isolates. If we focus on the three sites that repeatedly yielded isolates, we see a detection rate of 88.9% (8/9). This suggests that our ability to find enteroviruses circulating in Ibadan Nigeria can be optimized by focusing on these sites (drainage channels in Inalende [site 3], Gege [site 4] and Odo-Ona [site 5]). This study therefore highlights locations in Ibadan, Nigeria that can serve as ES sampling sites should the WHO program decide to include Ibadan in the surveillance program.

The results of this study show that CVB3, CVB6, E6, E7, E11, E12 and E13 (EV-Bs) were present in Ibadan metropolis during the study period. These EVs have been previously detected in ES and feacal samples in Nigeria (Adeniji and Faleye, 2014a; Oyero et al., 2014; Faleye and Adeniji, 2015a & b, Faleye et al., 2016) and beyond (Blomqvistet al., 2011, Tassin*et al*., 2013). Clinical presentations associated with these EV types range from AFP to neonatal systemic illness (Blomqvistet al., 2011, Tassin*et al*., 2013, Oyero et al., 2014). Currently, there is no case based surveillance system in place in the country to monitor the contribution of viruses, and particularly enteroviruses to these clinical conditions. Considering the eradication of poliovirus might now be within reach, the time seems right to put systems in place to monitor these viruses and probably investigate their contribution to different clinical manifestations in the country.

As crucial as case based surveillance is, we are not in any way suggesting that it should take precedence over ES. For example, it was found in this study that another lineage of E7 that was yet to be described was circulating in Ibadan, Nigeria (Figure 3). This lineage (Nigeria 2 within the sub-Saharan Africa cluster) had not been previously reported in association with any clinical manifestation and was not even described during ES in Lagos, Nigeria in 2012 (Faleye and Adeniji, 2015a) and 2013 (Faleye and Adeniji, 2015b). This is in consonance with the epidemiology of enteroviruses which specifies that majority of EV infections are sub-clinical and clinical manifestations only occur in one out of 100 to 250 infected people (Nathanson and Kew, 2010). This therefore highlights the capacity of ES to show EV strains that are not associated with clinical manifestations and might consequently not show up via case based surveillance studies.

The findings of this study also highlight the capacity of ES to function as an early warning system, signaling the EV strains that might be showing up in the clinic at a later time. For example, we found in Ibadan, Nigeria an E7 isolate that resurfaced in AFP cases over a year later and interestingly, in another geo-political zone of the country (Figure 3). This lineage (Nigeria within the Global Cluster) was detected in ES about a year before AFP associated cases showed up in the clinics. Hence, ES showed the silent circulation of this lineage before clinical associations were obvious. This capacity of ES gives us the insight necessary to acquire a rich and deep understanding of the dynamics and evolutionary trajectory of EV circulation. Knowledge that might be valuable for predictions and thereby facilitate the establishment of adequate preventive measures aimed to stop outbreaks early as is being done by the GPEI (WHO, 2004).

In this study, RD, HEp2C, MCF-7 and L20B cell lines were used for EVs isolation in a bid to increase the rate and diversity of EVs detected. No isolate was recovered on L20B cell line and consequently, no poliovirus was detected (Table 3). As expected, majority (68.8%) of isolates recovered on RD cell line were EV-Bs. This further confirms the EV-B bias of the cell line (Sadeuh-Mba et al., 2013; Adeniji and Faleye, 2014b; Faleye and Adeniji, 2015a & b). Though, HEp2C and MCF-7 have been documented to be EV-C bias (Sadeuh-Mba et al., 2013; Adeniji and Faleye, 2014b; Faleye and Adeniji, 2015b), the results of this study suggest that both cell lines might not be equivalent with respect to their susceptibility and permissiveness profile for EV-Cs. For example, while only two (50%) of the four enterovirus isolates recovered on HEp2C were completely identified, neither of the remaining two isolates was positive for the EV-C screen (Table 4). On the other hand, though only three (60%) of the five isolates recovered on MCF-7 were enteroviruses, two (66.7%) of the three enterovirus isolates were EV-Cs. More fascinating is the fact that one of the samples from site 4 yielded no isolate on RD and L20B cell lines but yielded CVB6 on HEp2C and a non-polio EV-C on MCF-7 (data not shown). It can be argued that this might just be a relic of the fact that two aliquots of the same ES concentrate might not contain the same virus diversity (Hovi et al., 2005; Adeniji and Faleye, 2014b). However, we feel otherwise because when isolates reported in Adeniji and Faleye (2014a) were passaged on HEp2C and MCF-7, all showed CPE on HEp2C but none showed CPE on MCF-7 (unpublished data). It therefore seems that though both appear to have some level of EV-C bias, HEp2C is less stringent in its selectivity for EV-Cs, while MCF-7 is more stringent in this respect.

As first shown in the emergence of several lineages of recombinant cVDPV2 in the country (Burns et al, 2013; 2014) with structural region of OPV2 origin and non-structural region of NPEV-C origin, and subsequently confirmed (Adeniji and Faleye, 2014b; Faleye and Adeniji 2015b), the results of this study further show that NPEV-C are present and circulating in Nigeria (Table 4 & Figure 4). It further reiterates the fact that the RD-L20B algorithm might be the major reason circulation and diversity of NPEV-Cs are under-reported in the country. Furthermore, the fact that the PV1 isolated in Cameroon in 2014 (Montmayeur et al., 2017) might have recombined with a non-structural genomic region similar to that of one of the NPEV-Cs described here suggests that likelihood of VDPV emergence is still real in Nigeria. Consequently, it is essential that effort be focused on ensuring there is herd immunity to all the poliovirus types especially now that elimination in the region seems within reach.

**Figure 4:**
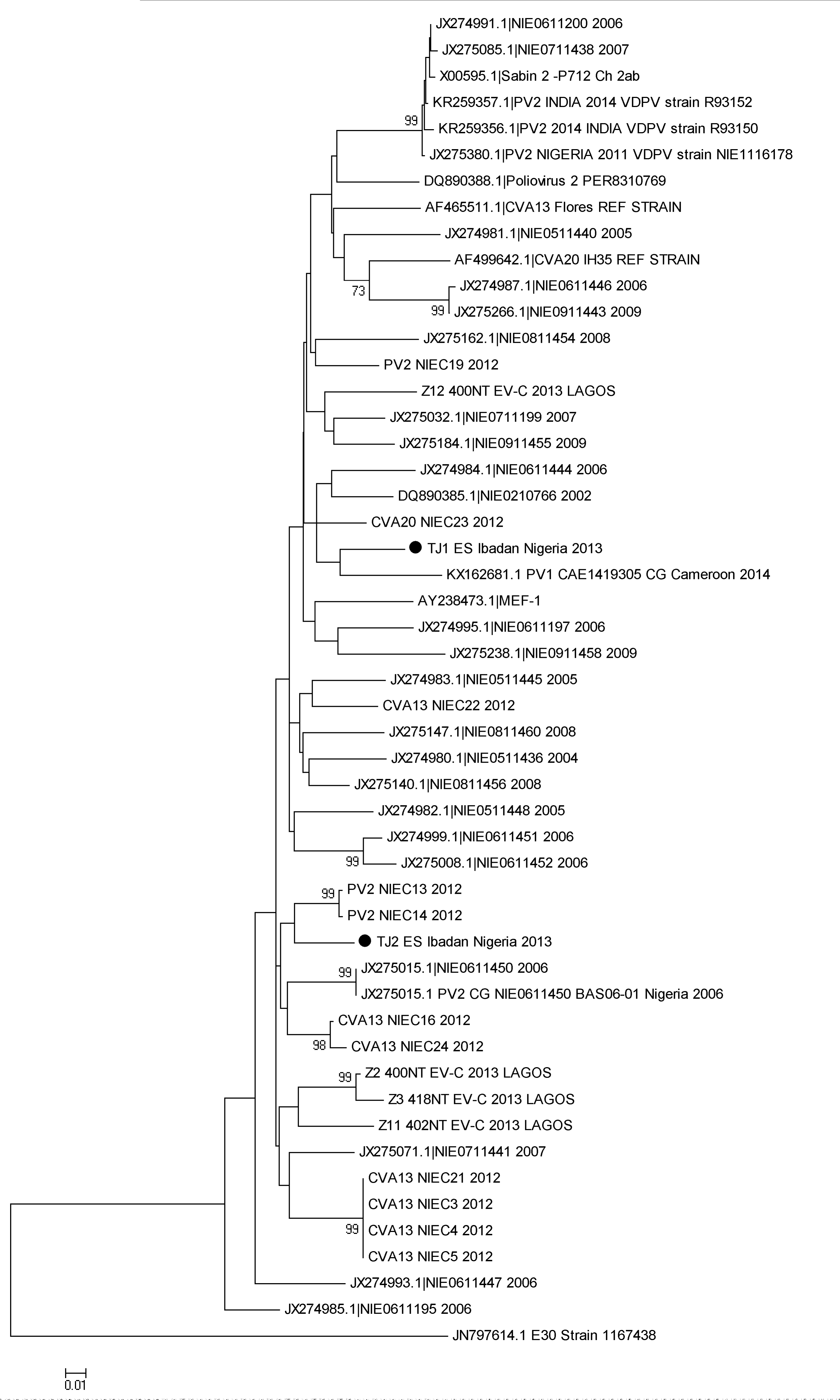
Phylogenetic relationship of isolates in the 3D/3^*I*^-UTR regions. The newly sequenced isolate and/or genomic region is indicated by black circles. Bootstrap values are indicated if > 50%.

Eight EV isolates that could not be identified were detected in this study. The reason these EVs could not be typed is not clear. It is possible that this might be as a result of mismatch of one or few bases near the end of the primer(s) or to more extensive nucleotide sequence divergence that resulted in the inability of the primers to amplify the VP1 gene of these isolates. However, in our lab, we have observed that using the primers AN32-AN35 (Nix et al., 2006) for cDNA synthesis instead of random hexamers (as used in this study) could result in successful amplification of the VP1 gene and consequently identification of previously unidentifiable isolates.

Finally, two isolates recovered on MCF-7 cell line were negative for the PanEnterovirus 5^l^-UTR screen. Alongside one other isolate we detected in Lagos, Nigeria in 2012 (Faleye and Adeniji, 2015a), as at the conclusion of this study, our lab had three isolates on MCF-7 cell line that appear not to be enteroviruses. This further emphasizes the need to define the range of viruses that MCF-7 cell line is both susceptible and permissive to.

## ACKNOWLEDGEMENTS

We thank the WHO National Polio Laboratory in Ibadan, Nigeria for providing the RD, HEp2C and L20B cell lines used in this study. We also thank Dr. James Ayorinde for the MCF-7 cell line used in this study.

## AUTHOR CONTRIBUTIONS

1. Study Design (All Authors)
2. Sample Collection, Laboratory and Data analysis (AOA and FTOC)
3. Wrote, revised, read and approved the final draft of the Manuscript (All Authors)

